# Mechanistic Insights into Human Cathelicidin LL-37 - Induced Activation of a Broadly Conserved Bacterial Signaling System

**DOI:** 10.1101/2025.05.06.652532

**Authors:** Samuel A. Adeleye, Abraham F. Mechesso, Arpita Mukherjee, Premal Shah, Guangshun Wang, Srujana S. Yadavalli

## Abstract

Antimicrobial peptides (AMPs) represent a promising class of therapeutics against bacterial pathogens. While their direct bactericidal mechanisms are well-characterized, how bacteria sense and respond to these peptides at sublethal concentrations remains poorly understood. Here, we investigate the activation of the *Escherichia coli* PhoQ-PhoP signaling system by the human antimicrobial peptide LL-37 and its derivatives (KR-12 and RI-10). We demonstrate that these peptides exhibit variable antimicrobial potency but surprisingly similar abilities to activate the PhoQ-PhoP pathway, indicating that signaling function is separable from bactericidal activity. Notably, sublethal concentrations of these peptides induce significant cell elongation, a phenotype dependent on PhoQ and mediated by the upregulation of QueE, which interferes with bacterial cell division. Contrary to the previous model suggesting peptides activate PhoQ passively by displacing its inhibitor MgrB, we observed enhanced cell elongation in *ΔmgrB* strains across all tested peptides, including RI-10, lacking antibacterial activity. Our findings suggest peptides actively stimulate PhoQ through a mechanism independent of MgrB dissociation, providing a more refined understanding of the peptide signaling through the PhoQ-PhoP system. These insights into bacterial adaptation mechanisms against host-derived peptides may guide the development of peptide therapeutics with enhanced efficacy against drug-resistant pathogens.

**Importance:** Antimicrobial peptides (AMPs) are promising alternatives to conventional antibiotics, yet how bacteria detect and respond to these host defense molecules remains poorly understood. This study investigates the interaction between bacterial sensing systems and AMPs at the genetic and molecular levels. Unlike other eukaryotes with multiple cathelicidin peptides, humans have only one cathelicidin that produces various active fragments through processing. Rather than creating multiple detectors, *E. coli* deploys an elegant solution, the PhoQ-PhoP signaling system that recognizes the conserved antimicrobial core region shared by all LL-37 fragments. We demonstrate that peptides directly stimulate PhoQ independent of their bactericidal activity, even in the absence of inhibitor MgrB, inducing cell elongation. These insights may inform the development of effective peptide-based therapeutics against drug-resistant pathogens.

## Introduction

When exposed to specific cationic antimicrobial peptides (AMPs) at sublethal concentrations, *Escherichia coli* activates the PhoQ/PhoP signal transduction pathway—a conserved two-component system crucial for stress adaptation in gram-negative bacteria (1, 2). The PhoQ/PhoP two-component signaling system is also an important pathway for adaptation and survival in conditions such as low Mg^2+^, mildly acidic pH, and hyperosmolarity in *E. coli* and closely related enterobacteria (3, 4). Activated PhoQ-PhoP regulate numerous genes, including those responsible for remodeling the outer membrane, activation of efflux pumps, and proteolytic degradation of the peptides that help bacteria resist cationic AMPs (5–8). Strong activation of the PhoQ-PhoP system is observed when cells are treated with a human platelet-derived AMP (C18G), causing the upregulation of QueE, an enzyme primarily known for its role in the queuosine biosynthetic pathway that modifies specific tRNAs(2). Remarkably, elevated QueE levels lead to the accumulation of this protein at the bacterial division septum, where it interacts with the cell division machinery. This interaction disrupts normal septation processes downstream of Z-ring formation, resulting in inhibited cell division and the development of filamentous bacterial morphology. This mechanism represents a sophisticated bacterial stress response that temporarily halts proliferation when facing host-antimicrobial challenges.

Host defense peptides (AMPs) are a key component of innate immunity, capable of rapidly eliminating invading pathogens (9, 10). Human cathelicidin, LL-37, is one of the best-studied AMPs (11–13) with a broad spectrum of activity against bacteria, fungi, parasites, viruses, and cancer cells (13–17). LL-37 is known to activate the PhoQ-PhoP signaling system and exert its antimicrobial action primarily by interacting with bacterial membranes, inserting itself into the lipid bilayer, and leading to membrane destabilization and the formation of pores (1, 18–20). This disruption results in the loss of membrane integrity, causing leakage of cellular contents and ultimately leading to bacterial cell death. Human cathelicidin LL-37 contains 37 amino acids and starts with a pair of leucine residues. Triple-resonance NMR structural studies established a long amphipathic helix, with the C-terminal tail of LL-37 (residues 33-37) disordered and not required for membrane-targeting mediated antimicrobial activity (21). Several labs used the LL-37 peptides from the central region for therapeutic development (13, 21, 22). KR-12 is a 12-amino-acid-long, minimal antibacterial peptide from human LL-37, which is toxic to bacteria but not human cells (21). It has been widely utilized in peptide engineering (23). To effectively develop AMPs as novel peptide therapeutics, it is imperative to study the bacterial response to these peptides, including the biochemical and regulatory pathways that underlie this process.

In natural environments, bacteria are far more likely to encounter sub-inhibitory AMP concentrations, conditions to which they have evolved dedicated sensing and resistance mechanisms, yet the physiological responses to these low-level exposures remain largely unknown (24, 25). Prior structural and biochemical studies established that cationic AMPs compete with Mg^2+^ ions for an acidic binding pocket in the PhoQ periplasmic sensor domain, directly triggering kinase activation (24, 25). Beyond divalent cations, peptides and other signals, a 47-amino-acid-long small membrane protein called MgrB interacts with and inhibits PhoQ by affecting its kinase activity (26, 27). Specific residues in the transmembrane and periplasmic regions of MgrB are important for the interaction with PhoQ (28). A recent model suggests that AMPs indirectly activate PhoQ by displacing MgrB and do not play an active role in stimulating PhoQ (29). There are two key mechanistic gaps arising from these models: Is the sensing of AMPs by PhoQ-PhoP and downstream gene activation dependent on peptide toxicity? And do AMPs directly stimulate PhoQ or act indirectly by relieving MgrB-mediated inhibition?

To gain deeper insights into the stress response mechanism to AMPs in *E. coli*, we utilized LL-37, its shortest anti-*E. coli* peptide KR-12, and an even shorter non-antibacterial peptide RI-10, resulting from the truncation of the N-terminal lysine and C-terminal arginine of KR-12. In particular, we first examined the antibacterial activity of LL-37 and its smaller fragments, KR-12 and RI-10, using different *E. coli* strains under physiologically relevant conditions. We determined the killing kinetics and characterized the peptides using membrane permeabilization and depolarization assays. Next, we investigated whether LL-37 and its fragments, at concentrations below MIC, are effective in activating the PhoQ/PhoP signaling system by analyzing both the changes in cell morphology and PhoP-dependent transcriptional reporter activity as readouts. Finally, based on our analysis of cells lacking the PhoQ inhibitor, MgrB, we propose a model of how peptides like LL-37 and its derivatives may interact with bacterial histidine kinase and affect bacterial physiology. Our study sheds light on the molecular mechanism of bacterial detection of the biologically relevant peptide LL-37, which is known to play a key role in human health via both antimicrobial and immune regulatory roles (9, 11–14). Our results reveal that the bacterial PhoQ-PhoP two-component system is able to detect a variety of LL-37 antimicrobial fragments in a specific manner by recognizing the consensus core antimicrobial region.

## Results

### Antibacterial activity of LL-37 and its derivatives against *E. coli*

To better delineate the antibacterial activity of LL-37, KR-12, and RI-10 against *E. coli*, we compared both their minimum inhibitory concentration (MIC) and minimum bactericidal concentration (MBC) against *E. coli* K-12 strain MG1655 (Tables 1, S1) under three different media conditions: Mueller-Hinton broth (MHB), which is a standard medium used for MIC assays (30), tryptic soy broth (TSB), and supplemented minimal A medium (MinA) (31), which are both more commonly used for bacterial growth. Gentamicin was included as a non-peptide antibiotic control. LL-37 exhibited the highest potency with MIC values between 2 and 8 μM, depending on the media used (Table S1). Interestingly, the KR-12 peptide, which is shorter than LL-37 by 25 amino acids, is also active against *E. coli*. The MIC and MBC values for KR-12 are about 2 to 4-fold higher than those obtained for LL-37. The values obtained in minimal medium were in a similar range as those obtained in other media (Table S1). The shortest LL-37 variant, RI-10, did not show antibacterial activity across different media conditions or after dilution of MHB to 10%. The data establish that the MIC values are generally consistent across different media; however, both MHB and TSB are undefined and complex, so we used the semi-defined supplemented minimal A medium as the growth medium from here on, as it is better suited than complex media for characterization of the physiological effects of these peptides, especially human LL-37. Helical wheel projections for LL-37, KR-12, and RI-10 indicate a hydrophobic side constituting a cluster of non-polar residues, distinct from the hydrophilic side, indicating a true helical potential (Figure 1). In contrast, KE-10 has low hydrophobicity with a negative <H> value, suggesting it is highly hydrophilic. Next, we investigated the time-dependent killing kinetics of LL-37 and KR-12 to further substantiate their bactericidal properties using a concentration corresponding to 2x MIC values (Figure S1). Since RI-10 does not display an MIC (>64 μM), we used 32 μM of this peptide to test the effect on bacterial killing. LL-37 at 2× MIC completely killed *E. coli* MG1655 cells grown to the exponential phase in 60 minutes, while KR-12 caused about a 4-log fold reduction in 2 hours. In contrast, RI-10 displayed no activity at the concentration tested. Together, these results show that full-length LL-37 and KR-12 exhibit antibacterial properties against *E. coli* but not RI-10.

**Figure 1.**
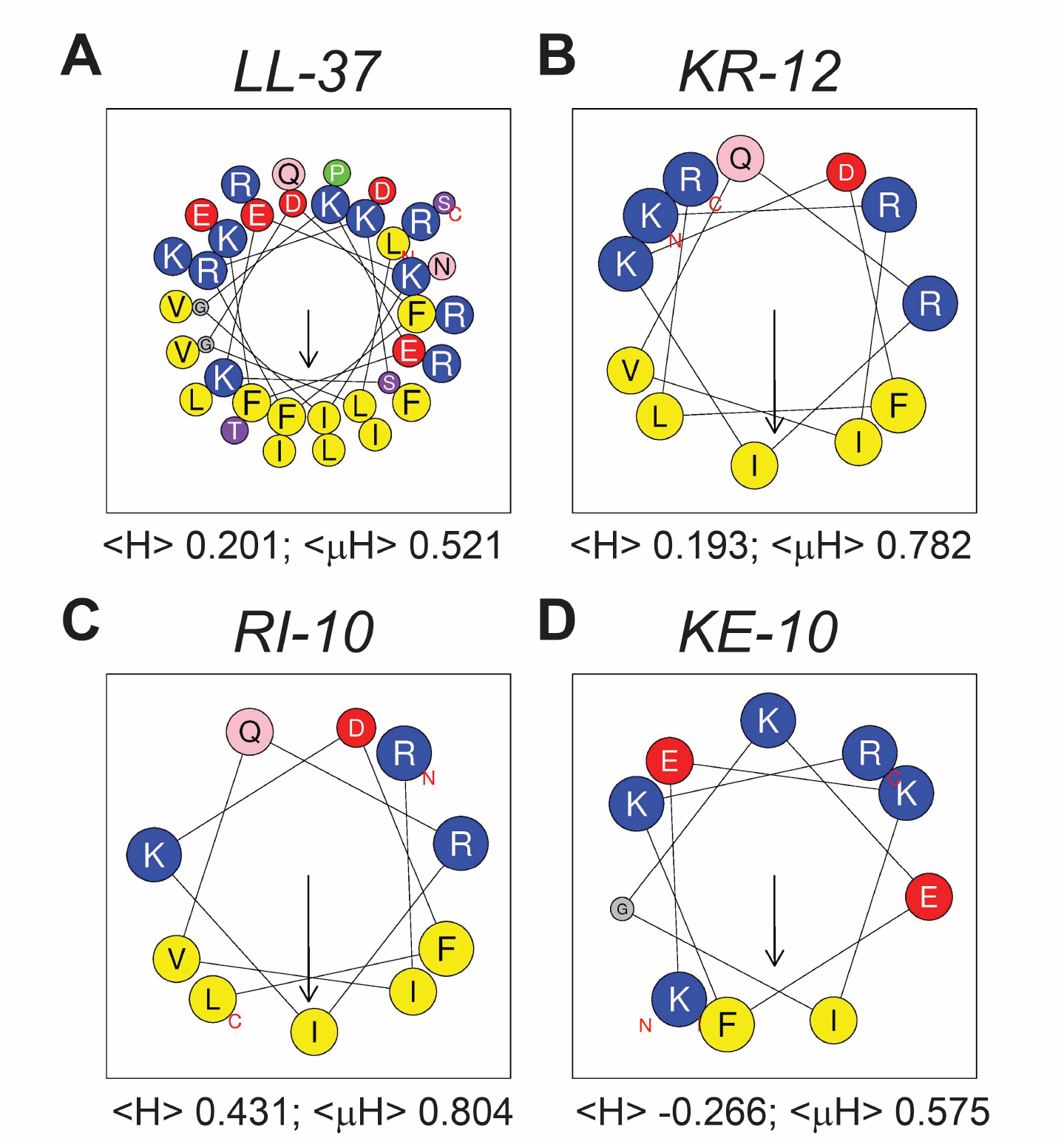
Helical wheel plots of human cathelicidin LL-37 and its fragments. (**A**) LL-37, (**B**) KR-12, (**C**) RI-10, and (**D**) KE-10. The arrow represents the hydrophobic moment vector and points to the center of the hydrophobic surface. The hydrophobicity <H> and hydrophobic moment <µH> are indicated underneath each peptide panel. Amino acids are color-coded as follows: blue: positively charged, red: negatively charged, yellow: hydrophobic, purple: S/T, pink: N/Q, grey: G, and green: P. Plots were generated using HeliQuest (46).

**Table 1.**
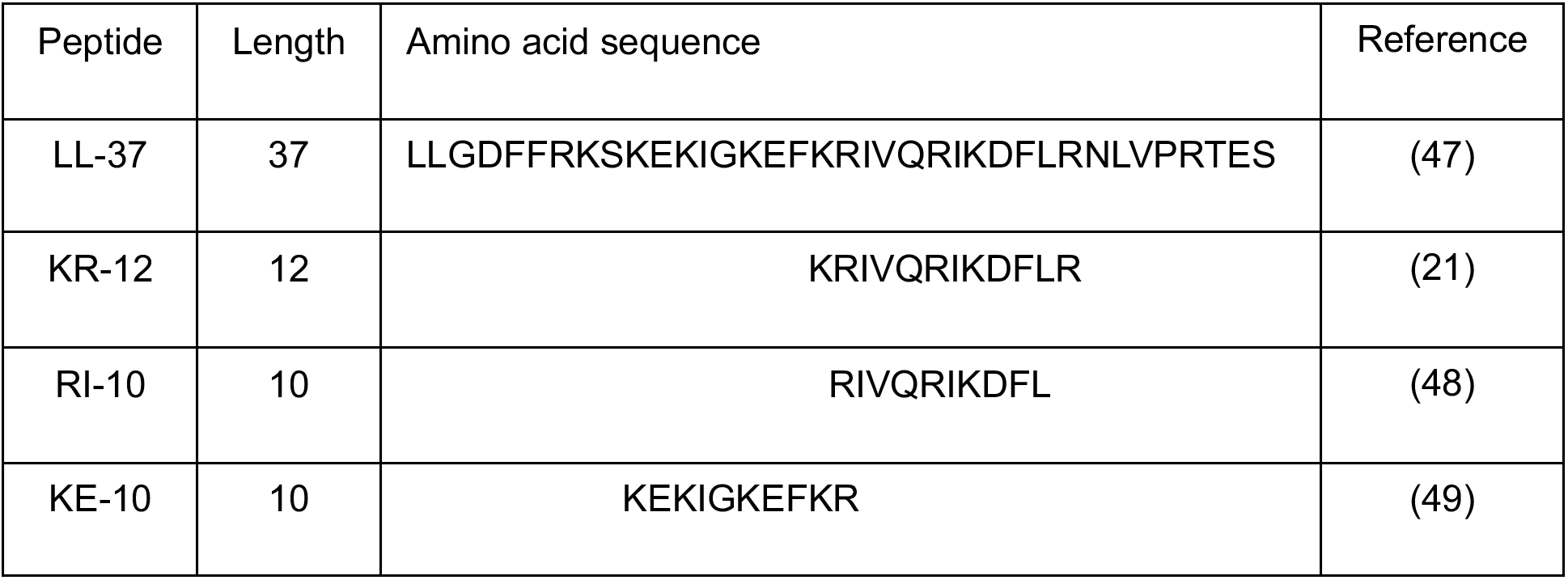
List of peptides used in the study.

### LL-37 and KR-12, but not RI-10, disrupt bacterial membranes

Given that LL-37 targets bacterial membranes, we wondered if the shorter peptide variants, KR-12 and RI-10, may retain and exhibit activity against *E. coli* membranes. To assess membrane disruption, we conducted membrane permeabilization and depolarization experiments. We analyzed membrane permeability by the uptake of a membrane-impermeable red dye, propidium iodide (PI), either in the presence of a peptide, 0.1% Triton-X (positive control) or PBS (untreated). Treatment with either LL-37 or KR-12 led to an increase in membrane permeability in a dose-dependent manner, although LL-37 was more effective (Figure S2A-C). To assay membrane depolarization, we used DiBAC_4_ (3), a probe for membrane potential, whose fluorescence increases upon entering depolarized cells. As expected, LL-37 depolarized membranes well with increasing concentration of the peptide (Figure S2D-F). Similarly, KR-12 also caused membrane depolarization but to a lower extent relative to LL-37. Consistent with the data from MIC assay and killing kinetics, RI-10 did not have a discernible effect on membrane permeability and depolarization. Together, these results suggest that KR-12, like LL-37, targets the bacterial membrane as its mode of action.

### Sub-MIC levels of LL-37 and KR-12 activate the PhoQ/PhoP signaling system and cause cell elongation

Cationic antimicrobial peptides, LL-37 and C18G activate the PhoQP pathway in *E. coli* and related bacteria (2, 28). In addition, *E. coli* cells elongate upon exposure to a sub-lethal dose of C18G, leading to filamentous growth (2, 29). This filamentation is associated with conditions that strongly activate PhoQ/PhoP. We asked if the truncated LL-37 derivatives retain the ability to activate the PhoQP system. We used a promoter-reporter strain containing P*_mgrB_*-*yfp* to measure the transcriptional activity of the *mgrB* gene, whose expression is PhoQP-dependent. In parallel, we observed the cells by phase contrast microscopy to test if the peptides induce any changes in *E. coli* cell morphology. Our analysis revealed that there is ∼3- to 4-fold increase in PhoQP-regulated transcriptional activity when cells are treated with a sub-MIC dose of LL-37 at 1.25 µM and KR-12 at 5 µM (Figure 2A). RI-10 showed weak activation of ∼2-fold higher than the untreated control. As the amino acid sequence of RI-10 (i.e., residues 19-28 of LL-37) is included in both KR-12 (residues 18-29) and LL-37, we also asked how a different LL-37 segment, KE-10 (residues 10-19), with a minimal sequence overlap with KR-12 and RI-10 (Table S1), behaves. Like RI-10, KE-10 showed no antibacterial activity against *E. coli* (MIC > 64 µM in Table S1). In contrast to RI-10, the KE-10 peptide showed negligible activation of the PhoQP system based on the transcription activity measurement of the *mgrB* gene (Figure 2A), implying that there is a requirement for a particular peptide sequence for this activation. None of the peptides, including LL-37 and KR-12, increased P*_mgrB_* reporter activity in *ΔphoQ* cells, confirming that activation is PhoQ-dependent (Figure 2B). Although LL-37 was long known to activate the PhoQP system, its impact on *E. coli* cell morphology, especially at concentrations below the MIC, has not been studied previously. Upon observation by phase-contrast microscopy, both LL-37 and KR-12-treated cells displayed elongation with varying cell lengths up to tens of microns (Figure 2C, E). In comparison to the untreated control, cells treated with LL-37 and KR-12 showed wide heterogeneity in cell size, and the average cell lengths increased by ∼1.6-fold with LL-37 and 1.3-fold with KR-12 (Figure 2C). RI-10 and KE-10-treated cells show a tighter distribution of cell lengths, with average values lower than those of LL-37 and KR-12 (Figure 2C, E). Although KE-10 treatment showed a statistically significant increase in average cell length, this difference is small and the cell sizes are quite uniform, lacking the heterogeneity observed in the case of LL-37 or KR-12, as evident from the violin plots. As expected, Δ*phoQ* cells did not elongate when treated with the test peptides (Figure 2D-E). Together, these data elucidate that LL-37, KR-12, and RI-10 are functional in stimulating the PhoQP-dependent promoter activity, but not KE-10. However, only full-length LL-37 and its minimal antibacterial peptide KR-12 are strong enough to cause elongation with increased heterogeneity in cell lengths similar to what was observed for the C18G peptide previously (2, 31), whereas two shorter LL-37 segments (RI-10 and KE-10), which lost antibacterial activity, were unable to.

**Figure 2.**
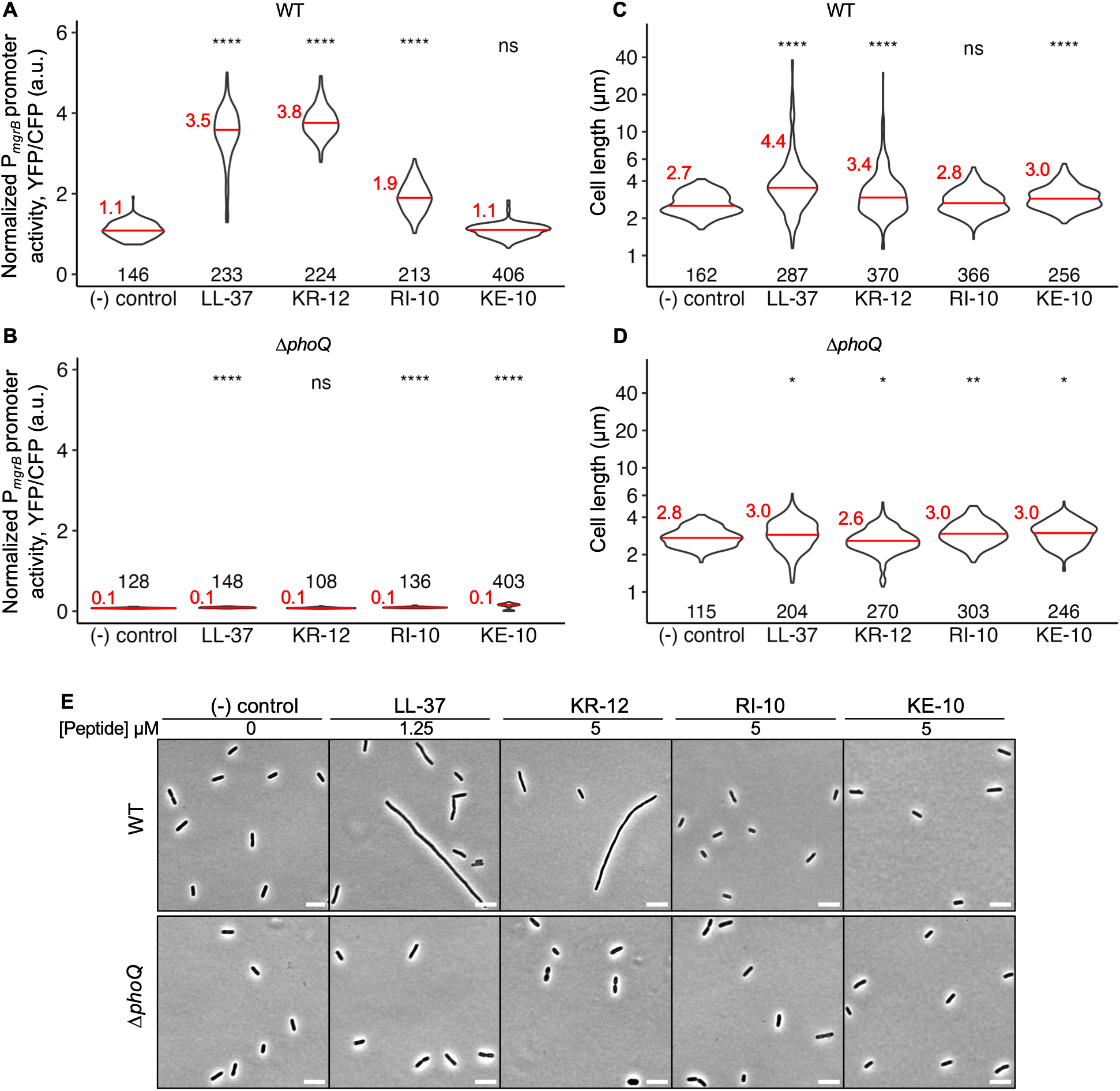
PhoQP-regulated transcription and cell morphology of *E. coli* cells treated with sub-MIC levels of antimicrobial peptides. **(A, B)** PhoQP-dependent promoter P*_mgrB_-yfp* reporter activity, a.u.: arbitrary units, **(C, D)** Quantification of cell lengths, and, **(E)** Representative phase contrast micrographs, scale bar = 5 µm, of *E. coli* MG1655 (TIM92) and *ΔphoQ* (TIM100) cells, respectively, grown in the presence of sub-MIC levels of antimicrobial peptides LL-37 (1.25 µM), KR-12 (5 µM), RI-10 (5 µM), and KE-10 (5 µM) in supplemented MinA with 0.1 mM MgSO_4_. All data and images shown in the above panels A-C are obtained from at least three biological replicates; mean and median values are shown in red text and red bars, respectively. The number of cells analyzed is indicated for each sample. Significance is determined by the Shapiro-Wilk test: ****P ≤ 0.0001, *P ≤ 0.05, and “ns” = P > 0.05.

### Cell elongation caused by sub-MIC levels of peptides is dependent on QueE

In previous work, sub-MIC levels of C18G are shown to cause filamentation in wild-type *E. coli* cells, which is mediated by increased expression of a PhoP-dependent gene called *queE*, encoding a protein with dual activities in the inhibition of cell division and the synthesis of queuosine tRNA modification (2, 32). We asked if the morphological phenotype observed here for cells exposed to LL-37 and KR-12 is also mediated by QueE or through an independent mechanism. Using a *queE* deletion strain, we carried out microscopy and measured cell lengths and transcriptional reporter activity. It is worth noting that Δ*queE* cells grown in the presence of these peptides still display an increase in PhoQP-regulated transcriptional reporter activity similar to the wild-type cells (Figures 3A, 2A). Δ*queE* cells exposed to peptides did not show a difference in the average cell length relative to the controls (no peptide, KE-10, Figure 3B), and Δ*queE* suppressed the varied cell elongation phenotype observed when cells were treated with either LL-37 or KR-12 (Figures 3B-C). Overall, filamentation and heterogeneity in cell size observed upon treatment with LL-37 and KR-12 peptides are QueE-dependent.

**Figure 3.**
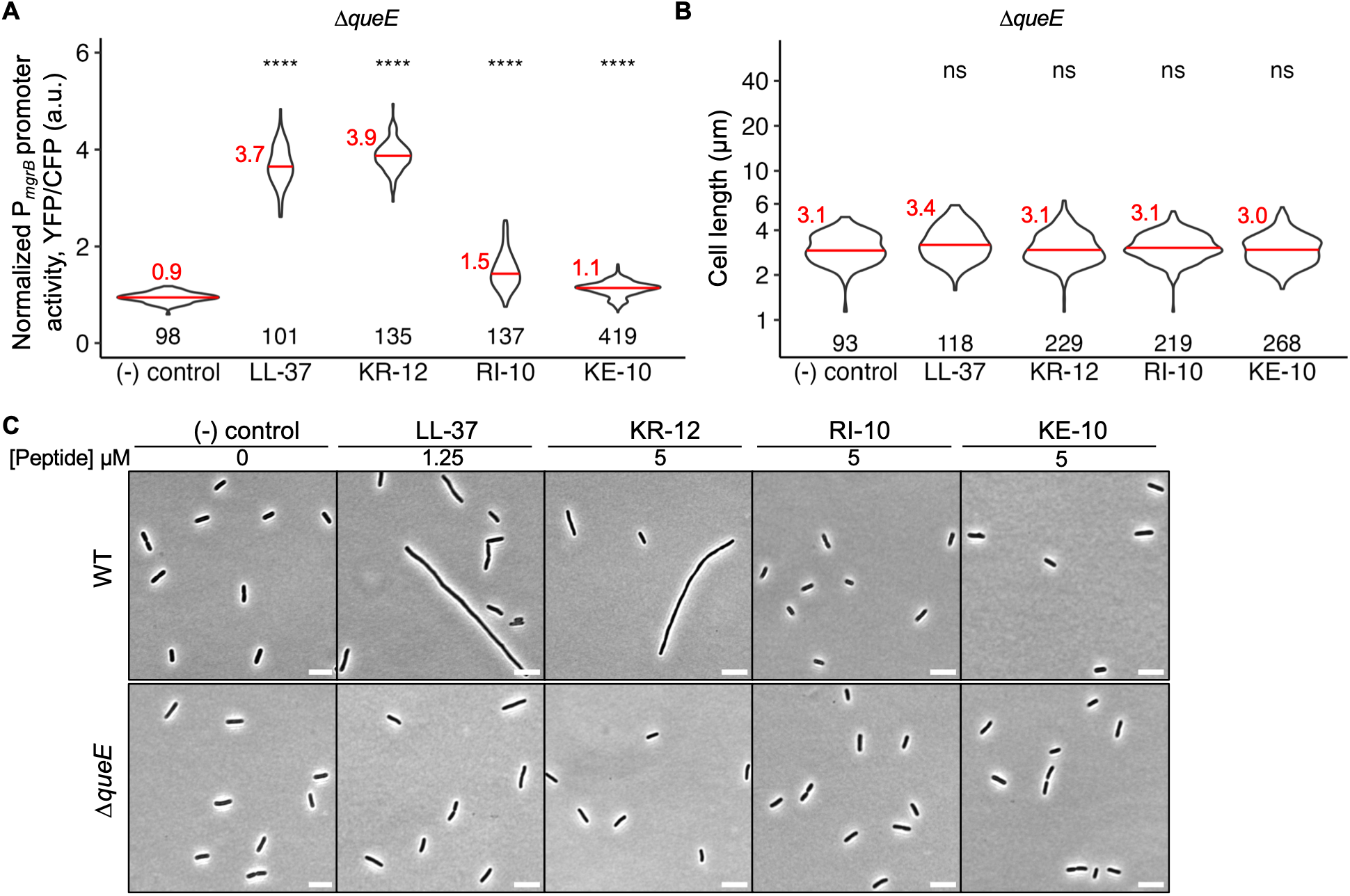
PhoQP-regulated transcription and cell morphology of *ΔqueE* cells treated with sub-MIC levels of antimicrobial peptides. **(A)** PhoQP-dependent promoter P*_mgrB_-yfp* reporter activity, a.u.: arbitrary units, **(B)** Quantification of cell lengths, and **(C)** Representative phase contrast micrographs, scale bar = 5 µm, of Δ*queE* (SAM31) cells grown in the presence of sub-MIC levels of the antimicrobial peptides LL-37 (1.25 µM), KR-12 (5 µM), RI-10 (5 µM), and KE-10 (5 µM) in supplemented MinA with 0.1 mM MgSO_4_. WT (TIM92) cell cultures were included as a control for cell length and promoter activity measurements, which yielded reproducible results as represented in the WT data shown in Figure 2. All data and images shown in the above panels A-E are obtained from at least three biological replicates; mean and median values are shown in red text and red bars, respectively. The number of cells analyzed is indicated for each sample. Significance is determined by the Shapiro-Wilk test: ****P ≤ 0.0001, ***P ≤ 0.001, *P ≤ 0.05, and “ns” = P > 0.05.

### Deletion of PhoQ inhibitor, MgrB, increases cell responsiveness to LL-37 and its derivatives

*E. coli* cells carrying an *mgrB* deletion show an overall, robust increase in PhoQP-controlled gene expression as PhoQ is uninhibited (23). These cells experience higher stimulation of PhoQ under magnesium-limiting conditions, similar to the strong activation of this system observed in wild-type cells grown in the presence of C18G at intermediate magnesium concentrations (24). Consequently, Δ*mgrB* cells grow as long filaments, and this phenotype is dependent on upregulation of QueE expression by PhoQ/PhoP (26). It is proposed that MgrB mediates the sensing of cationic AMPs by PhoQ at intermediate concentrations of magnesium, therefore, LL-37, KR-12, and RI-10 may activate PhoQ only when MgrB is present. In this case, we expect Δ*mgrB* cells to show no significant change in either the PhoP-dependent transcriptional reporter activity or cell morphology when treated with peptides vs. no treatment. We examined how LL-37 and its derivatives influence the activity of the PhoQP system and cell lengths in a Δ*mgrB* strain. As expected, in the absence of peptides, Δ*mgrB* cells showed an increase in P*_mgrB_* transcription relative to the wild-type cells under the same growth conditions (Figures S3, 4A). When the AMPs were added to Δ*mgrB* cells, there was a modest increase of ∼1.4-fold in the PhoQP-regulated transcriptional reporter activity for LL-37, as observed previously using the same media conditions carrying 0.1 mM Mg^2+^ (Figures 4A, S3) (29). The addition of peptides KR-12, RI-10, and KE-10 did not lead to a substantial increase in reporter activity, suggesting that PhoP-mediated transcription is not sensitive in response to the two peptides in cells lacking MgrB. This result is consistent with a previous study where Δ*mgrB* cells did not show an increase in PhoP-regulated transcription in the presence of another AMP, C18G (27).

**Figure 4.**
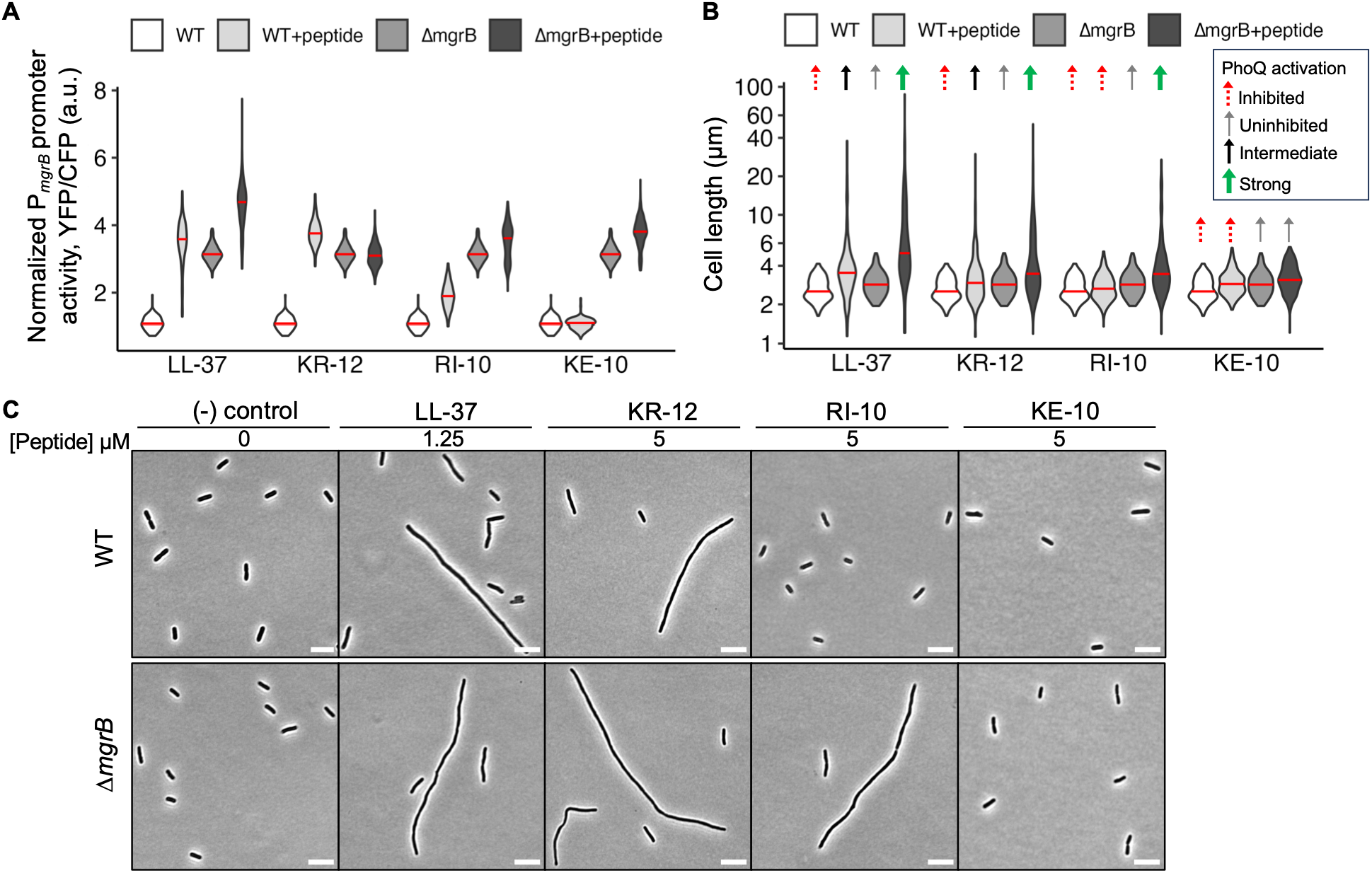
PhoQP-regulated transcription and cell morphology of *ΔmgrB* cells treated with sub-MIC levels of antimicrobial peptides. **(A)** PhoQP-dependent promoter P*_mgrB_-yfp* reporter activity in *ΔmgrB* (AML20) cells relative to WT *E. coli* (TIM92), a.u.: arbitrary units, **(B)** Comparison of cell lengths in *ΔmgrB* (AML20) cells with those of WT *E. coli* (TIM92), and **(C)** Representative phase contrast micrographs of WT *E. coli* (TIM92) and *ΔmgrB* (AML20) cells, scale bar = 5 µm. Cells were grown in the presence of sub-MIC levels of the antimicrobial peptides LL-37 (1.25 µM), KR-12 (5 µM), RI-10 (5 µM), and KE-10 (5 µM) in supplemented MinA with 0.1 mM MgSO_4_. Arrows in panel (B) indicate the PhoQ activation status. WT (TIM92) cell cultures were included as a control for cell length and promoter activity measurements, which yielded reproducible results as represented in the WT data shown in Figure 2. All data and images shown in the above panels A-C are obtained from at least three biological replicates; median values are shown in red bars. See Figure S3 in the Supplementary Material for the P*_mgrB_* promoter activity and cell length data for *ΔmgrB* (AML20) cells, including the number of cells analyzed per sample and statistical analysis.

To investigate this further, we analyzed the morphology of Δ*mgrB* cells in the presence of peptides, wherein cell elongation is PhoQ-dependent and a direct function of strong PhoQP activation. As expected, we noticed filamentation in Δ*mgrB* cells treated with LL-37 and KR-12, but only short, typical cells in the KE-10-treated or no-peptide control cells, under this growth condition with 0.1 mM Mg^2+^ (Figures 4B-C, S3). It was intriguing, however, to see that RI-10-treated Δ*mgrB* cells also displayed elongation, suggesting that RI-10 may interact more effectively with PhoQ to activate it, in the absence of MgrB. In other words, loss of the inhibitor MgrB appears to unmask the effect of the RI-10 peptide on PhoQ stimulation, resulting in an evident cell elongation phenotype. Interestingly, cell size quantification revealed greater heterogeneity and higher average cell lengths across all three peptides for Δ*mgrB* of 6-8 μm (Figure 4B-C, S3). The mean cell lengths for LL-37, KR-12, and RI-10-treated Δ*mgrB* cells were higher than the corresponding values obtained for the wild type by 2.5-, 2.3-, and 1.9-fold, respectively; however, there is no difference in cell length in the KE-10-treated cells (Figures 4B-C, S3), indicating a central role of the RI-10-containing LL-37 segments in activation in the absence of *mgrB*.

It is known that high stimulation of PhoQ from exposure to LL-37 results in differential regulation of specific PhoP-regulated promoters, including that of *mgrB* (33). Also, upon activation by LL-37, a mutated *mgrB-t7a* promoter (with a T7A substitution in the PhoP-binding site) shows a higher fold change in transcriptional activity relative to the wild-type *mgrB* promoter. To test whether this modified promoter that is highly sensitive to antimicrobial peptides through activation of the PhoQ-PhoP pathway, may elicit a strong morphological response, we analyzed wild-type and Δ*mgrB* strains carrying the *mgrB-t7a* promoter. We observed a significant increase in cell size heterogeneity in wild-type cells treated with LL-37 and KR-12, and Δ*mgrB* cells treated with LL-37, KR-12, and RI-10 showed an exacerbated cell elongation phenotype with increased cell lengths in the strain carrying the modified *mgrB-t7a* promoter (Figure S4) relative to the unmodified promoter sequence for *mgrB* (Figure 4). Notably, both the WT and Δ*mgrB* cells did not display an increase in promoter activity or cell length when treated with control peptide KE-10 or in the absence of a peptide (Figure S4). If the exacerbated cell elongation observed in peptide-treated Δ*mgrB* cells is indeed due to a PhoQP-dependent mechanism that is mediated by QueE, we should see this phenotype suppressed upon deletion of *queE*. Consistent with our hypothesis, Δ*mgrB* Δ*queE* cells treated with LL-37, KR-12, and RI-10 did not show an increase in PhoQ-dependent transcriptional reporter activity and displayed a loss of filamentation and a decrease in cell size heterogeneity (Figure S5). Together, these data suggest that LL-37 and its derivatives bind and actively stimulate PhoQ. In the absence of MgrB, PhoQ is uninhibited, and these peptides further enhance PhoQ activation. Importantly, cell elongation serves as a sensitive phenotype for peptide-mediated PhoQ stimulation in *E. coli*. Wild-type cells exposed to sub-MIC peptide concentrations exhibit moderate elongation, reflecting intermediate PhoQ activity, while *ΔmgrB* cells display significantly increased cell lengths, indicating robust PhoQ activation (Figure 4B).

## Discussion

Despite extensive investigation, fundamental questions remain about the molecular mechanisms of peptide-mediated signal transduction by the PhoQ-PhoP system. Moreover, the effects of PhoPQ activation by cationic AMPs in cells and the physiological importance of these interactions have not been elucidated. We address two critical questions in this work: (i) Can peptides lacking potent antimicrobial activity still interact with PhoQ and act as a signal for this pathway? Using LL-37 and derivatives with variable MICs and antibacterial activities against *E. coli* cells, we show that the signaling function of peptides through the PhoQ-PhoP system is separable from their antimicrobial activity (Figure 5A), suggesting that bacteria may use this pathway primarily as a detection mechanism for host environments rather than simply as a response to membrane damage. (ii) Do the peptides actively stimulate PhoQ, or do they exert their effect on PhoQ passively by displacing the inhibitor protein, MgrB? Many studies on peptide-induced signaling of PhoQ measure the activity of peptides using PhoP-dependent transcriptional reporters as a proxy. In agreement with the previous studies, upon exposure to peptides, we do not observe a significant increase in PhoQP-driven transcriptional reporter activity in *ΔmgrB* cells relative to the wild type. It is possible that the derepression of PhoP-regulated gene expression in the absence of MgrB could lead to lower sensitivity of the reporter signal, masking small increments in the amplitude. While such reporters are incredibly useful in determining the activation status of this pathway and measuring the transcription levels, we have a second, sensitive readout – a cell elongation or filamentation phenotype in response to peptide stress in *E. coli*. Based on our microscopy and cell length measurements, we find that cells elongate significantly upon exposure to sublethal concentrations of peptides, LL-37 and KR-12, even though the promoter reporters do not show a significant change in their activities. The extent of cell elongation may differ in response to specific peptides, as we note that filamentation induced by an AMP, C18G, in wild-type *E. coli* cells is quite robust, with an average cell length of > 20 μm (30). Upon deletion of MgrB, the average cell lengths increased further, and even RI-10-treated cells displayed cell elongation; this observation suggests that AMPs may activate PhoQ directly in addition to causing MgrB dissociation (32). Our findings elucidate PhoQ-dependent effects of these peptides and support a model where peptides play an active and direct role in PhoQ activation (Figure 5B). At high Mg²⁺ (millimolar concentrations), Mg²⁺ occupies the acidic patch of the PhoQ periplasmic domain and outcompetes AMPs, and PhoQ remains repressed (1). At physiological Mg²⁺ (∼0.1-1 mM), cationic peptides stimulate PhoQ, while MgrB limits peptide-induced signaling, yielding intermediate pathway output; deleting *mgrB* removes this constraint, so peptide exposure produces maximal activation. At low Mg²⁺ (0-100 µM), cells experience slow growth, and PhoQ is activated independently of peptide (33); addition of AMPs imposes further stress through membrane damage.

**Figure 5.**
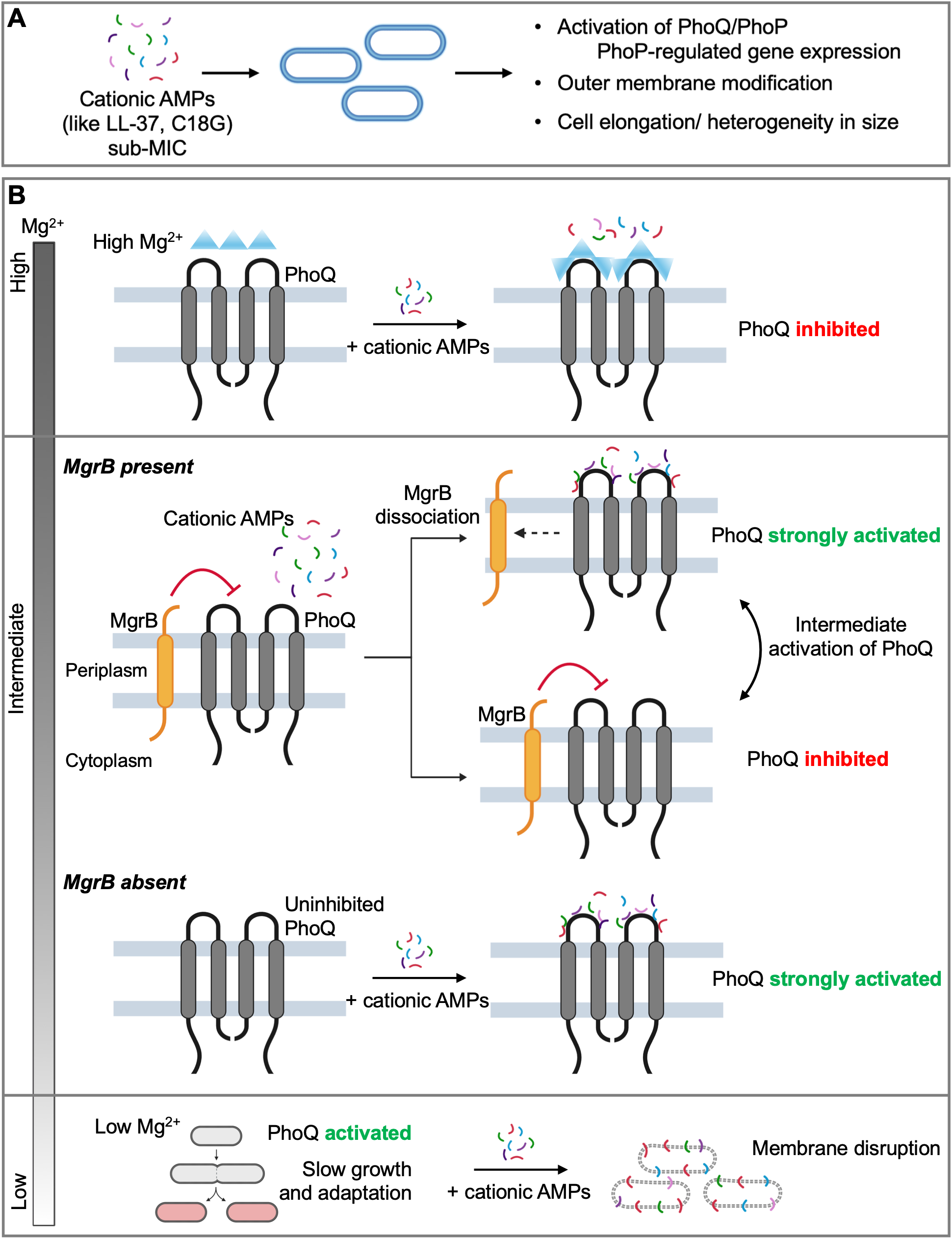
Schematic overview of peptide stress response through the *E. coli* PhoQ-PhoP signaling system. **(A)** At high concentrations, antimicrobial peptides (AMPs) affect membrane integrity and cause damage, leading to cell death. At sub-MIC levels, peptides are sensed by the signaling systems, such as PhoQ/PhoP, which control gene expression under peptide stress, leading to physiological changes, including cell elongation and modification of the outer cell membrane. **(B)** At high concentrations of magnesium (Mg^2+^, ≥1 mM), AMPs are outcompeted by Mg^2+^ ions and PhoQ is inhibited. At physiological (intermediate Mg^2+^, ∼0.1-1 mM) magnesium concentrations, PhoQ is strongly stimulated by specific cationic peptides, and some PhoQ molecules are inhibited by the small protein regulator MgrB, leading to intermediate PhoQ activation. In the absence of MgrB, PhoQ is not inhibited, and when cells are exposed to peptides, PhoQ is strongly activated, resulting in maximal activation of this signaling system. At low concentrations of Mg^2+^ (0-100 μM), bacterial cell growth is slow, accompanied by high activation of PhoQ, and addition of AMPs imposes additional stress through membrane disruption.

Our understanding of what characteristics of AMPs are required to activate the PhoP-PhoQ system is still evolving. A previous report found a correlation between the extent of positive charge, hydrophobicity, and amphipathicity of an AMP and its ability to activate PhoQ-PhoP (34). Intriguingly, the authors discovered that even a peptide with minimal positive charge (+1) but containing histidines was highly potent at activating the system, suggesting the existence of an activation mechanism more complex than thought. A later study with 117 AMPs from the Antimicrobial Peptide Database (34) showed that PhoQ is activated by a diverse set of AMPs, including the human cathelicidin LL-37 and non-cathelicidin peptides that carry either an α-helical or a combination of α-helical/β-sheet structural regions (35). In our work, we see that LL-37 derivatives induce PhoQ-dependent morphological changes, and KR-12, the smallest antibacterial peptide, is also the smallest peptide sequence for PhoPQ induction. Further, RI-10, but not KE-10, became responsive upon the deletion of *mgrB*. Interestingly, LL-25, the N-terminal segment of LL-37, which does not kill bacteria under normal conditions (35), does not activate PhoPQ based on the recent human AMPs screen study, although it contains a partial sequence of KR-12 (34). LL-25 can be obtained by extending the sequence of KE-10 from both ends (Tables 1, S2). Together, the data from the current study suggest that part of the LL-37 amino acid sequence that overlaps with both KR-12 and RI-10 is likely mediating the interaction with PhoQ. More broadly, our findings imply that the bacterial molecular detection machinery is elegantly tuned to specifically target the core antimicrobial region of human antimicrobial peptide LL-37 but not fragments outside this region. When broadly viewed from the host-pathogen interaction at the molecular level, bacterial detection of the core antimicrobial region of human cathelicidin LL-37 (35–39) is remarkably efficient since this region is commonly shared by all the active fragments of LL-37 naturally produced in human skin, including KS-27, KR-20, KS-30, RK-31, and LL-29 (Table S2) (36). Indeed, all these human cathelicidin fragments responded to the PhoPQ system based on a recent screening of human AMPs registered in the APD (34). Collectively, these findings underscore that the bacterial PhoQ-PhoP system likely senses host defense peptides based on evolutionarily conserved structural and chemical properties independent of their antimicrobial activity, allowing bacteria to mount preemptive responses rather than simply react to membrane damage.

Finally, the dramatic rise of antimicrobial resistance is a major public health concern. AMPs and other peptide derivatives are relatively small, typically comprised of around 10 to 50 amino acids (34), and constitute a key component of host innate immunity for defense against microbial pathogens (36, 37). These cationic peptides are relatively easy to produce, have good thermal stability and water solubility, and exhibit a broad spectrum of activity against bacteria and fungi, including clinical isolates of drug-resistant *E. coli* (Table S5), making them promising candidates for drug development (38–42). Studying the bacterial responses to antimicrobial peptides in general provides us with a fundamental understanding of how pathogens can sense, adapt, and develop tolerance to these host defense molecules and guide the strategic design of novel therapeutics with enhanced efficacy to overcome the growing antibiotic resistance problem.

## Methods

### Strains

Details of the strains used in this study are provided in Supplementary Table S3. AML20, JNC21, NRS6, SAM31, TIM92, TIM100, and TIM166 are *E. coli* K-12 MG1655 derivatives. Additionally, clinical strains of *E. coli*, E423-17, E416-17, and ATCC strain 25922 were used for specific assays as indicated. Strain NRS6 was made as follows: KEIO strain JW1815 (carrying *ΔmgrB*::*kan^R^*) was infected with phage P1_vir_, and the resulting phage lysate was used to transduce the TIM166 strain (33), respectively, using P1_vir_ (42).

### Media, reagents, and growth conditions

Routine bacterial growth on solid agar was performed using LB Miller medium (IBI Scientific) containing 1.5% bacteriological grade agar (VWR Life Sciences) at 37°C. Liquid cultures were grown at 37°C with aeration in either LB Miller medium or minimal A medium (MinA) (42) (K_2_HPO_4_ (10.5 g l^-1^), KH_2_PO_4_ (4.5 g l^-1^), (NH_4_)_2_SO_4_ (1.0 g l^-1^), Na_3_ citrate.2H_2_O (0.5 g l^-1^), supplemented with 0.1% casamino acids, 0.2% glucose and 1 mM MgSO_4_ unless otherwise indicated. Throughout this study, supplemented MinA, as indicated above, will be referred to as MinA or minimal medium. In addition, Mueller-Hinton broth (MHB, BD Difco DF0757-17-6) and tryptic soy broth (TSB, Wards® 470227-500) were used for minimum inhibitory/bactericidal concentration (MIC/MBC) assays. Peptides and/or AMPs used in the study are indicated in Table 1.

For the measurement of cell length and transcriptional reporter activity, overnight cultures of TIM92 (WT), TIM100 (*ΔphoQ*), SAM31 (*ΔqueE*), AML20 (*ΔmgrB*), JNC21 (*ΔmgrB ΔqueE*), NRS6 (WT-2), and TIM166 (*ΔmgrB*-2) were grown in 3 ml supplemented MinA containing 1 mM MgSO_4_ in borosilicate glass culture tubes with aeration. Saturated cultures were backdiluted 1:500 into 5 ml supplemented MinA with 0.1 mM Mg^2+^ and grown for 2 hours in a shaker at 300 rpm. Each culture was split into two tubes, with one tube designated as a no-peptide control and the second tube where a test peptide was added. LL-37, KR-12, RI-10, and KE-10 were added to a final concentration of 1.25 µM, 5 µM, 5 µM, and 5 µM, respectively. These cultures were further grown for 4 hours at 37°C in a roller drum at 60 rpm. TIM92 (WT) cultures were always included as a control for cell length and promoter activity measurements for each of the other strains under different peptide treatments. At least 3 independent biological replicates were included per strain for each treatment. For a given strain, the promoter activity and cell length measurements were highly reproducible and consistent across replicates from different days of experimentation.

### Microscopy and image analysis

Cell morphology was observed and single-cell fluorescence to measure transcriptional reporter activity was performed using phase contrast and fluorescence microscopy as previously described (43). 3 µl of the cells were immobilized on a 1% agarose pad and observed using a Nikon TiE fluorescent microscope with a TI2-S-HU attachable mechanical stage and a perfect focus system (PFS). The images were captured using a Teledyne Photometrics Prime 95B sCMOS camera with 1x1 binning. YFP and CFP fluorescence was imaged using a 100 ms exposure time at 20% intensity. Image acquisition was done using Metamorph software version 7.10.3.279 (Molecular Devices, Sunnyvale, CA). The background fluorescence was determined using MG1655 cells grown under the same conditions. The average cell fluorescence was quantified using ImageJ (44) and the MicrobeJ plugin (45). Data from independent replicates were plotted using R.

### Minimum inhibitory/bactericidal concentration (MIC/MBC) testing

The antibacterial activity of the peptides against *E. coli* MG1655, E423-17, E416-17, and 25922 strains was evaluated using a previously established protocol (45). Briefly, a peptide concentration gradient with twofold dilution was prepared in the 96-well polystyrene microplate at 10 μl per well. Bacteria were grown to the exponential phase in MHB, TSB, or supplemented MinA. It was then diluted to ∼10^5^ CFU/ml in MHB, TSB, or supplemented MinA with 0.1 mM MgSO_4_, and partitioned into the 96-well microplate at 90 μl per well. After overnight incubation at 37°C, readings were taken at 600 nm using a ChroMate 4300 Microplate Reader at (GMI, Ramsey, MN). The minimum inhibitory concentration (MIC), represented as ranges, was determined for wells without bacterial growth. For minimum bactericidal concentration (MBC) determination, aliquots of 30 μl from growth-absent wells were plated on MH agar plates, and the lowest peptide concentration with no isolated colony was considered the MBC. Results were expressed in µM.

### Membrane permeabilization

The peptide ladder was prepared as described for the antibacterial assay in a black COSTAR 96-well plate. Propidium iodide (PI) (MP Biomedicals, Solon, OH) was prepared in the dark and dissolved in DMSO (Thermo Fisher Scientific, NY) to 20 mM. This PI stock solution was further diluted to 1 mM with water and 2 μl of 1 mM PI was added to each well. An overnight culture of *E. coli* K12 MG1655 was inoculated into 1xMinA supplemented with 0.2% glucose, 0.1% casamino acids, 0.1 mM MgSO_4_, and grown to exponential phase. It was then diluted to OD_600_ of 0.11 in the same medium containing peptides at the indicated concentrations, and 88 μl was added to each well. The plate was incubated at 37°C with continuous shaking in a FLUOstar Omega (BMG LABTECH, NC) microplate reader. The sample was read every 5 minutes for 24 cycles with excitation and emission wavelengths of 584 nm and 620 nm, respectively.

### Killing kinetics

*E. coli* K12 MG1655 was grown to the exponential phase (OD_600_ ∼ 0.3) in 1xMinA supplemented with 0.2% glucose, 0.1% casamino acids, 1 mM MgSO_4_. The culture was diluted to OD_600_ 0.001 (∼10^5^ CFU/ml) in 1xMinA supplemented with 0.2% glucose, 0.1% casamino acids, 0.1mM MgSO_4_. One milliliter of bacterial suspension was then mixed with LL-37 (4 μM, 2× MIC), KR-12 (8 μM, 2× MIC) or RI-10 (32 μM) and incubated at 37°C. After 15, 30, 60, 90, and 120 minutes of incubation, 50 μl was taken and serially diluted with 1×PBS. Then, 50 μl of the diluted suspension was plated on MH agar plates. The plates were incubated overnight at 37°C for bacterial CFU determination.

### Membrane depolarization

An overnight culture of *E. coli* K12 MG1655 was inoculated into 1xMinA supplemented with 0.2% glucose, 0.1% casamino acids, 1 mM MgSO_4_, and grown to the exponential phase. Bacteria were washed with 1×PBS, resuspended in twice the volume of 1×PBS containing 25 mM glucose, and incubated at 37°C for 15 minutes. Then, 500 nM (final concentration) of the DiBAC4 (3) bis-(1,3-dibutylbarbituric acid) trimethine oxonol (ANASPEC, CA) was added and vortexed gently. Aliquots of 90 µl of the energized bacteria solution were loaded to the 96 well plates (Corning COSTAR, AZ) and placed in a FLUOstar Omega microplate reader (BMG LABTECH, NC). Fluorescence was read for 15 min at excitation and emission wavelengths of 485 nm and 520 nm, respectively. Then, 10 µl of peptide solutions were added, and fluorescence readings were recorded for 45 min. Triton X-100 (0.1%) was used as a positive control.

### Statistical analysis

In the datasets for cell lengths and promoter reporter activities, data were assessed for normal distribution using the Shapiro-Wilk test. For YFP/CFP fluorescence ratios, 15 of 25 groups departed significantly from normality (p < 0.05), with Shapiro-Wilk W statistics ranging from 0.929 to 0.995 across groups. For cell length, data were analyzed on the log_10_ scale, consistent with the data representation and the biological interpretation of fold changes in cell length; on this scale, 19 of 25 groups remained non-normal, with W statistics as low as 0.885 (p < 10^-16^ for *ΔmgrB* + RI-10). Given that the data are not distributed normally, and the distributional shape differed substantially across groups, Wilcoxon rank-sum tests (Mann-Whitney U) were applied consistently across all comparisons. The figures reflect Holm-corrected significance annotations throughout. All raw p-values and adjusted p-values are reported in Supplementary Table S4.

## Data, materials, and resource availability

The main manuscript and its supporting files include all data generated or analyzed during this study. Requests for strains and plasmids generated in this study should be directed to and will be fulfilled by the lead contact, Srujana S. Yadavalli. Further information and requests for resources and reagents should be directed to and will be fulfilled by Srujana S. Yadavalli and Guangshun Wang.

## Acknowledgments

The authors thank Dr. Mark Goulian for sharing transcriptional reporter strains for *mgrB*, Dr. Nitish Sharma for preparing *ΔmgrB* strains carrying the reporters, Dr. Manuela Roggiani for critical feedback on the manuscript, and members of the Yadavalli and Wang labs for helpful discussions. **Funding:** S.S.Y. is supported by the National Institutes of Health - National Institute of General Medical Sciences (NIH-NIGMS) ESI-MIRA R35 GM147566 and institutional start-up funds from Rutgers. A.M. is supported by the Waksman Institute Busch Predoctoral fellowship (2024-25) at Rutgers. G.W. is supported by the NIH/NIAID R56 AI175209. The funders did not play any role in the study design, data collection and analysis, decision to publish, or preparation of the manuscript.

## Author contributions

S.S.Y. and G.W. conceived the experiments. S.A.A., A.F.M., A.M., G.W., and S.S.Y. contributed to the planning of the experiments, designing the methodology, and data analysis. S.A.A., A.F.M., and A.M. performed the experiments. P.S. contributed to the generation of violin plots and performed statistical analysis. All authors contributed to the writing, review, and editing of the manuscript.

