## Supplemental Material for "Mechanistic Insights into Human Cathelicidin LL-37 - Induced Activation of a Broadly Conserved Bacterial Signaling System"

### Supporting Information

#### Tables

**Table S1. Minimum inhibitory concentration (MIC) and minimum bactericidal concentration (MBC) values for LL-37 and its derivatives against *E. coli* K-12 MG1655**. MIC values (μM) of the antimicrobial peptide LL-37 and its derivatives KR-12, RI-10, and KE-10 against *E. coli* K-12 MG1655 in supplemented minimal medium (MinA), Mueller-Hinton broth (MHB), and tryptic soy broth (TSB). The corresponding MBC values (μM) are shown within brackets. Gentamicin is included as a bactericidal antibiotic control. Data represent values obtained from at least three independent biological replicates.

| **Strain** | **Media** | **LL-37** | **KR-12** | **RI-10** | **KE-10** | **Gentamicin** |
|---|---|---|---|---|---|---|
| *E. coli* K-12 MG1655 | MinA | 2 [2] | 8-16 [16] | >64 [>64] | >64 [>64] | 0.5-1 [1] |
| | MHB | 8 [8] | 16-32 [32] | >64 [>64] | >64 [>64] | 0.5-1 [1] |
| | TSB | 8 [16] | 16 [32] | >64 [>64] | >64 [>64] | 1-2 [2] |
| *E. coli* K-12 MG1655 Δ*mgrB* | MinA | 2 [2] | 8 [16] | >64 [>64] | >64 [>64] | - |
| | MHB | 8 [8] | 8-16 [16] | >64 [>64] | >64 [>64] | - |
| | TSB | 8-16 [32] | 16 [32] | >64 [>64] | >64 [>64] | - |

**Table S2**. List of human LL-37 fragments that share the key recognition sequence for bacterial sensing (bolded).

| Name | Peptide amino acid sequence |
|---|---|
| LL-37 | LLGDFFRKSKEKIGKEF**KRIVQRIKDFLR**NLVPRTES |
| LL-23 | LLGDFFRKSKEKIGKEFKRIVQR |
| LL-29 | LLGDFFRKSKEKIGKEF**KRIVQRIKDFLR** |
| KR-20 | **KRIVQRIKDFLR**NLVPRTES |
| KS-27 | KSKEKIGKEF**KRIVQRIKDFLR**NLVPR |
| KS-30 | KSKEKIGKEF**KRIVQRIKDFLR**NLVPRTES |
| RK-31 | RKSKEKIGKEF**KRIVQRIKDFLR**NLVPRTES |

**Table S3.** List of strains used in the study.

| Strain | Relevant genotype | Reference/source |
|---|---|---|
| *E. coli* K12 MG1655 | $\lambda^-$ *rph-1* | *E. coli* Genetic Stock Center #7740 |
| *E. coli* E423-17 | Clinical antibiotic-resistant isolate | Wang lab, University of Nebraska Medical Center |
| *E. coli* E416-17 | Clinical antibiotic-resistant isolate | Wang lab, University of Nebraska Medical Center |
| *E. coli* 25922 | O6 isolate, reference strain for antibiotic susceptibility testing | ATCC |
| AML20 | MG1655 *ΔmgrB* $\lambda_{att}$ ($P_{mgrB}$-yfp) $HK_{att}$ ($P_{tetA}$-*cfp*) | (1) |
| JNC21 | MG1655 *ΔmgrB ΔqueE*::kan $\lambda_{att}$ ($P_{mgrB}$-*yfp*) $HK_{att}$ ($P_{tetA}$-*cfp*). $kan^r$ | (2) |
| JW1815 | BW25113 *ΔmgrB750::kan* | (3) |
| NRS6 | MG1655 *ΔmgrB::kan* *ΔphoQ* $\lambda_{att}$::($P_{mgrBt7a}$-*yfp*) $HK_{att}$::($P_{tetA}$-*cfp)* | This work |
| SAM31 | MG1655 *ΔqueE* $\lambda_{att}$ ($P_{mgrB}$-*yfp*) $HK_{att}$ ($P_{tetA}$-*cfp*) | (2) |
| TIM92 | MG1655 $\lambda_{att}$ ($P_{mgrB}$-yfp) $HK_{att}$ ($P_{tetA}$-*cfp*) | (4) |
| TIM100 | MG1655 *ΔphoQ* $\lambda_{att}$::($P_{mgrB}$-*yfp*) $HK_{att}$::($P_{tetA}$-*cfp)* | (4) |
| TIM166 | MG1655 *ΔphoQ* $\lambda_{att}$::($P_{mgrBt7a}$-*yfp*) $HK_{att}$::($P_{tetA}$-*cfp)* | (4) |

**Table S4. Statistical analysis of promoter activity and cell length datasets.** Raw p-values and adjusted p-values for all the data related to promoter activity and cell length measurements are reported below. The abbreviations are as follows: AMP, antimicrobial peptide; n_treatment = number of samples for a given peptide treatment; n_control = number of samples for no peptide control; W = Shapiro-Wilk test statistic for normal distribution (closer to 1 indicates normal distribution); p_raw = unadjusted p-value; p_holm = Holm-adjusted p-value; sig = significance based on p_holm.

| **Promoter activity (YFP: CFP) data** | | | | | | | |
|---|---|---|---|---|---|---|---|
| **AMP** | **n_treatment** | **n_control** | **W** | **p_raw** | **p_holm** | **sig** | **Strain** |
| LL-37 | 233 | 146 | 33960 | 5.89E-60 | 2.36E-59 | **** | TIM92 (WT) |
| KR-12 | 224 | 146 | 32704 | 1.85E-59 | 5.55E-59 | **** | TIM92 (WT) |
| RI-10 | 213 | 146 | 30318 | 8.91E-53 | 1.78E-52 | **** | TIM92 (WT) |
| KE-10 | 406 | 146 | 28186 | 0.37982056 | 0.37982056 | ns | TIM92 (WT) |
| LL-37 | 255 | 240 | 58833 | 1.69E-70 | 6.77E-70 | **** | AML20 (*ΔmgrB*) |
| KR-12 | 186 | 240 | 20341 | 0.11645633 | 0.11645633 | ns | AML20 (*ΔmgrB*) |
| RI-10 | 213 | 240 | 36420 | 5.78E-15 | 1.16E-14 | **** | AML20 (*ΔmgrB*) |
| KE-10 | 343 | 240 | 73212 | 1.03E-57 | 3.10E-57 | **** | AML20 (*ΔmgrB*) |
| LL-37 | 221 | 273 | 26099 | 0.00993709 | 0.02981127 | * | JNC21 (*ΔmgrB ΔqueE*) |
| KR-12 | 211 | 273 | 27893 | 0.55178295 | 0.55178295 | ns | JNC21 (*ΔmgrB ΔqueE*) |
| RI-10 | 216 | 273 | 32866 | 0.02931805 | 0.0586361 | ns | JNC21 (*ΔmgrB ΔqueE*) |
| KE-10 | 328 | 273 | 67724 | 2.51E-27 | 1.00E-26 | **** | JNC21 (*ΔmgrB ΔqueE*) |
| LL-37 | 101 | 98 | 9898 | 3.80E-34 | 1.14E-33 | **** | SAM31 (*ΔqueE*) |
| KR-12 | 135 | 98 | 13230 | 9.09E-39 | 3.64E-38 | **** | SAM31 (*ΔqueE*) |
| RI-10 | 137 | 98 | 12813 | 1.69E-32 | 3.38E-32 | **** | SAM31 (*ΔqueE*) |
| KE-10 | 419 | 98 | 34587 | 4.70E-26 | 4.70E-26 | **** | SAM31 (*ΔqueE*) |
| LL-37 | 148 | 128 | 14854 | 4.02E-16 | 8.04E-16 | **** | TIM100 (*ΔphoQ*) |
| KR-12 | 108 | 128 | 7620 | 0.17572905 | 0.17572905 | ns | TIM100 (*ΔphoQ*) |
| RI-10 | 136 | 128 | 14511 | 7.61E-21 | 3.04E-20 | **** | TIM100 (*ΔphoQ*) |
| KE-10 | 403 | 128 | 38386 | 8.24E-17 | 2.47E-16 | **** | TIM100 (*ΔphoQ*) |
| LL-37 | 154 | 417 | 60490 | 3.57E-59 | 1.07E-58 | **** | TIM166 (WT-2) |
| KR-12 | 154 | 417 | 64218 | 3.18E-75 | 1.27E-74 | **** | TIM166 (WT-2) |
| RI-10 | 418 | 417 | 126130 | 4.83E-29 | 4.83E-29 | **** | TIM166 (WT-2) |

| | | | | | | | |
|---|---|---|---|---|---|---|---|
| KE-10 | 171 | 417 | 5954 | 9.47E-57 | 1.89E-56 | **** | TIM166 (WT-2) |
| LL-37 | 69 | 241 | 15834 | 2.26E-30 | 6.78E-30 | **** | NRS6 (*ΔmgrB*-2) |
| KR-12 | 69 | 241 | 15576 | 1.95E-28 | 3.90E-28 | **** | NRS6 (*ΔmgrB*-2) |
| RI-10 | 84 | 241 | 18896 | 2.71E-32 | 1.08E-31 | **** | NRS6 (*ΔmgrB*-2) |
| KE-10 | 127 | 241 | 16760 | 0.13340228 | 0.13340228 | ns | NRS6 (*ΔmgrB*-2) |
| **Cell length data** | | | | | | | |
| **AMP** | **n_treatment** | **n_control** | **W** | **p_raw** | **p_holm** | **sig** | **Strain** |
| LL-37 | 287 | 162 | 35183 | 1.58E-19 | 6.32E-19 | **** | TIM92 (WT) |
| KR-12 | 370 | 162 | 38122 | 5.86E-07 | 1.52E-06 | **** | TIM92 (WT) |
| RI-10 | 366 | 162 | 32067 | 0.1343489 | 0.1343489 | ns | TIM92 (WT) |
| KE-10 | 256 | 162 | 26782 | 5.06E-07 | 1.52E-06 | **** | TIM92 (WT) |
| LL-37 | 168 | 188 | 27380 | 6.19E-33 | 2.48E-32 | **** | AML20 (*ΔmgrB*) |
| KR-12 | 352 | 188 | 45284 | 1.66E-12 | 3.32E-12 | **** | AML20 (*ΔmgrB*) |
| RI-10 | 399 | 188 | 51753 | 1.08E-13 | 3.23E-13 | **** | AML20 (*ΔmgrB*) |
| KE-10 | 196 | 188 | 22368 | 2.87E-04 | 2.87E-04 | *** | AML20 (*ΔmgrB*) |
| LL-37 | 249 | 263 | 40944 | 9.54E-07 | 2.86E-06 | **** | JNC21 (*ΔmgrB ΔqueE*) |
| KR-12 | 482 | 263 | 68982.5 | 0.04609891 | 0.04609891 | * | JNC21 (*ΔmgrB ΔqueE*) |
| RI-10 | 505 | 263 | 73049.5 | 0.02281511 | 0.04563022 | * | JNC21 (*ΔmgrB ΔqueE*) |
| KE-10 | 210 | 263 | 35973 | 1.53E-08 | 6.11E-08 | **** | JNC21 (*ΔmgrB ΔqueE*) |
| LL-37 | 118 | 93 | 6392 | 0.03995356 | 0.15981426 | ns | SAM31 (*ΔqueE*) |
| KR-12 | 229 | 93 | 10608.5 | 0.95839228 | 1 | ns | SAM31 (*ΔqueE*) |
| RI-10 | 219 | 93 | 10803.5 | 0.39534544 | 1 | ns | SAM31 (*ΔqueE*) |
| KE-10 | 268 | 93 | 11704 | 0.38233889 | 1 | ns | SAM31 (*ΔqueE*) |
| LL-37 | 204 | 115 | 13415 | 0.03319468 | 0.03319468 | * | TIM100 (*ΔphoQ*) |
| KR-12 | 270 | 115 | 12769.5 | 0.00583869 | 0.01518505 | * | TIM100 (*ΔphoQ*) |
| RI-10 | 303 | 115 | 20926.5 | 0.00149182 | 0.00596726 | ** | TIM100 (*ΔphoQ*) |
| KE-10 | 246 | 115 | 16735 | 0.00506168 | 0.01518505 | * | TIM100 (*ΔphoQ*) |
| LL-37 | 137 | 135 | 14524 | 4.16E-16 | 1.67E-15 | **** | TIM166 (WT-2) |
| KR-12 | 97 | 135 | 10495 | 4.97E-15 | 1.49E-14 | **** | TIM166 (WT-2) |
| RI-10 | 130 | 135 | 11077 | 2.24E-04 | 4.49E-04 | *** | TIM166 (WT-2) |
| KE-10 | 155 | 135 | 9643 | 0.25025571 | 0.25025571 | ns | TIM166 (WT-2) |
| LL-37 | 83 | 200 | 14285 | 1.32E-21 | 5.30E-21 | **** | NRS6 (*ΔmgrB*-2) |
| KR-12 | 59 | 200 | 9693 | 6.36E-14 | 1.91E-13 | **** | NRS6 (*ΔmgrB*-2) |
| RI-10 | 133 | 200 | 18987 | 3.87E-11 | 7.75E-11 | **** | NRS6 (*ΔmgrB*-2) |
| KE-10 | 99 | 200 | 6176 | 1.21E-07 | 1.21E-07 | **** | NRS6 (*ΔmgrB*-2) |

**Table S5. Minimum Inhibitory Concentration (MIC) and Minimum Bactericidal Concentration (MBC) values for LL-37 and its derivatives against clinical isolates of *E. coli***. MIC values (μM) of antimicrobial peptide LL-37 and its derivatives KR-12 and RI-10 against clinical, antibiotic-resistant isolates of *E. coli* in supplemented minimal medium (MinA), Mueller-Hinton broth (MHB), and tryptic soy broth (TSB). The corresponding MBC values (μM) are shown within brackets. Data represent values obtained from at least three independent biological replicates.

| ***E. coli* strains** | **Media** | **LL-37** | **KR-12** | **RI-10** | **Gentamicin** |
|---|---|---|---|---|---|
| | MinA | 2-4 [4] | 8 [8] | >64 [>64] | 2 [2] |
| *E.coli* E423-17 | MHB | 4-8 [8] | 32 [32] | >64 [>64] | 1 [1] |
| | TSB | 4-8 [8] | 32 [>32] | >64 [>64] | 2 [4] |
| | MinA | 4 [4] | 4-8 [8] | >64 [>64] | 2 [2] |
| *E.coli* E416-17 | MHB | 8 [8] | 16 [16] | >64 [>64] | 1 [2] |
| | TSB | 8-16 [32] | 16-32 [32] | >64 [>64] | 4 [4] |
| | MinA | 2-4 [8] | 1-2 [16] | >64 [>64] | 2 [4] |
| *E.coli* 25922 | MHB | 8 [8] | 16 [32] | >64 [>64] | 1 [2] |
| | TSB | 8 [32] | 32 [>32] | >64 [>64] | 2-4 [4] |

**Figures**

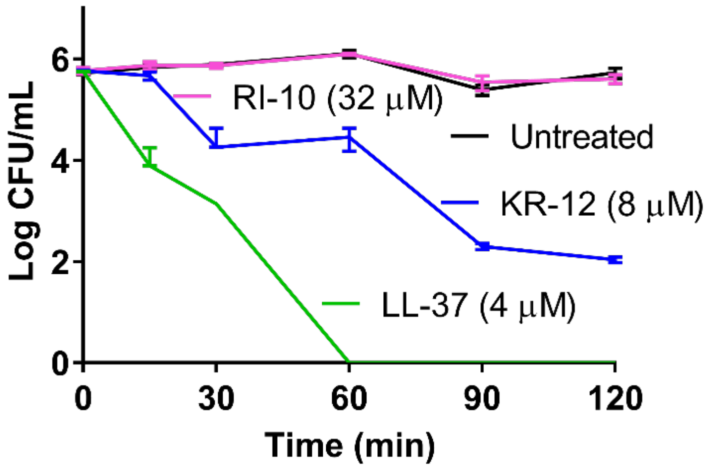

**Figure S1**. **Killing kinetics LL-37, KR-12, and RI-10 against *E. coli* K-12 MG1655.** Cells were treated with peptides at indicated concentrations in supplemented MinA medium with 0.1 mM $MgSO_4$. Data are represented as mean ± range for two biological replicates.

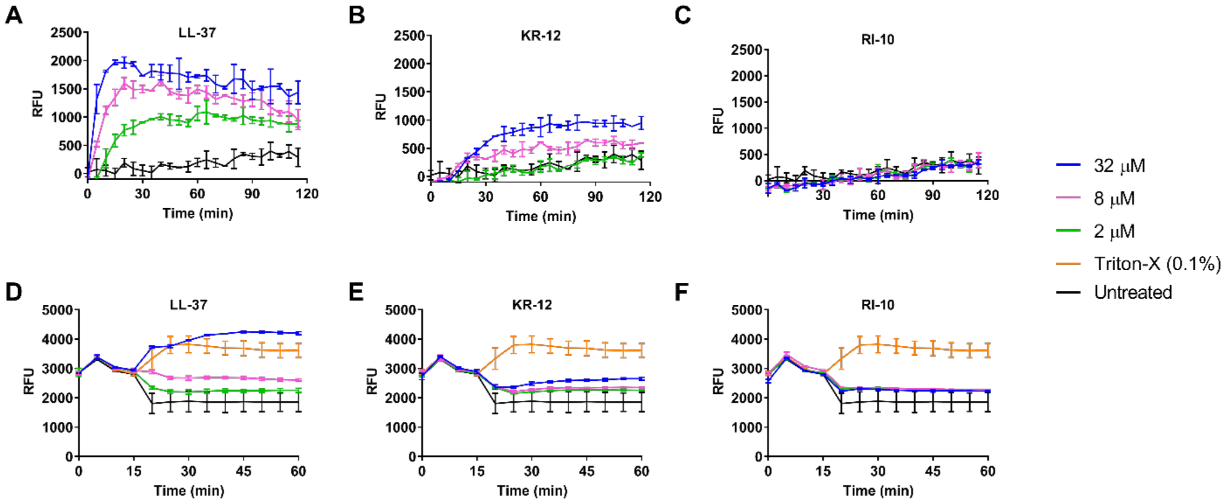

**Figure S2. Membrane permeabilization (A-C) and depolarization (D-F) activity of a series of LL-37, KR-12, and RI-10 in *E. coli* K-12 MG1655.** The assay was performed in supplemented MinA medium with 0.1 mM $MgSO_4$. RFU: Relative fluorescence intensity. Data are represented as mean ± range for two independent replicates.

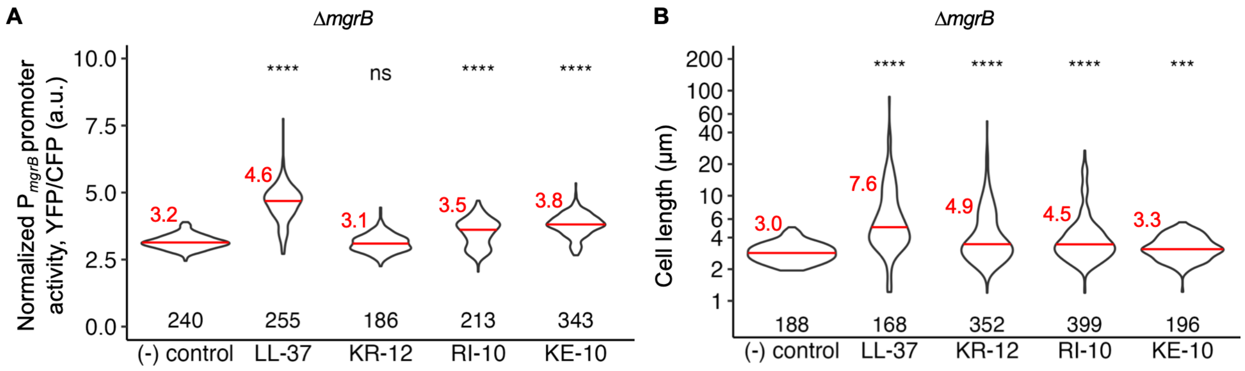

**Figure S3. PhoQP-regulated transcription and cell morphology of Δ*mgrB* cells treated with sub-MIC levels of antimicrobial peptides. (A)** PhoQP-dependent promoter $P_{mgrB}$-*yfp* reporter activity in Δ*mgrB* (AML20) cells, a.u.: arbitrary units, **(B)** Quantification of cell lengths of Δ*mgrB* (AML20) cells, grown in the presence of sub-MIC levels of the antimicrobial peptides LL-37 (1.25 µM), KR-12 (5 µM), RI-10 (5 µM), and KE-10 (5 µM) in supplemented MinA with 0.1 mM $MgSO_4$. Data were obtained from three biological replicates; mean and median values are shown in red text and red bars, respectively. The number of cells analyzed is indicated for each sample. Significance is determined by Shapiro-Wilk test: ****P ≤ 0.0001, ***P ≤ 0.001, **P ≤ 0.01, *P ≤ 0.05, and “ns” = P > 0.05.

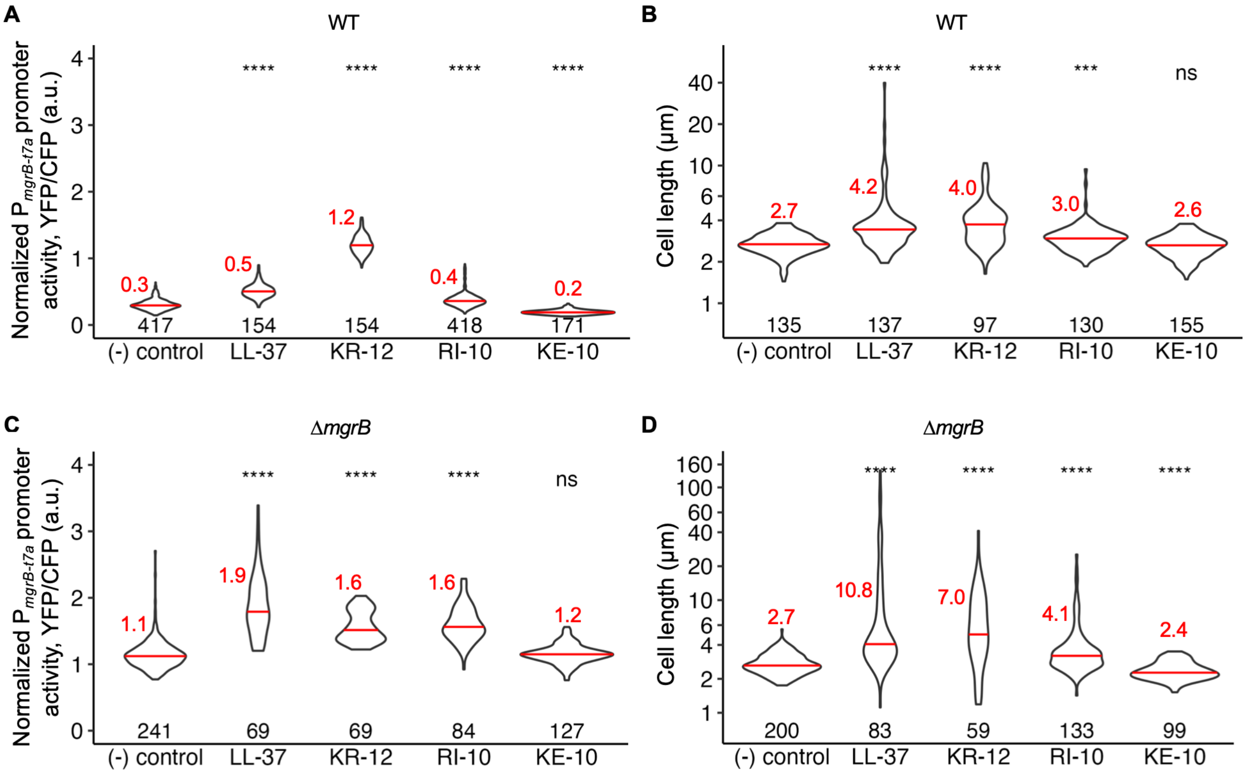

**Figure S4. PhoQP-regulated transcription and cell lengths in *mgrB-t7a* reporter cells treated with sub-MIC levels of antimicrobial peptides. (A, C)** PhoQP-dependent promoter reporter activity in wild-type (TIM166) or Δ*mgrB* (NRS6) cells, and **(B, D)** Quantification of cell lengths of wild-type or Δ*mgrB* (NRS6) cells, grown in the presence of sub-MIC levels of the antimicrobial peptides LL-37 (1.25 μM), KR-12 (5 μM), RI-10 (5 μM), and KE-10 (5 μM) in supplemented MinA with 0.1 mM $MgSO_4$. Data were obtained from three biological replicates; mean and median values are shown in red text and red bars, respectively. The number of cells analyzed is indicated for each sample. Significance is determined by Shapiro-Wilk test: ****P ≤ 0.0001, ***P ≤ 0.001, **P ≤ 0.01, *P ≤ 0.05, and “ns” = P > 0.05.

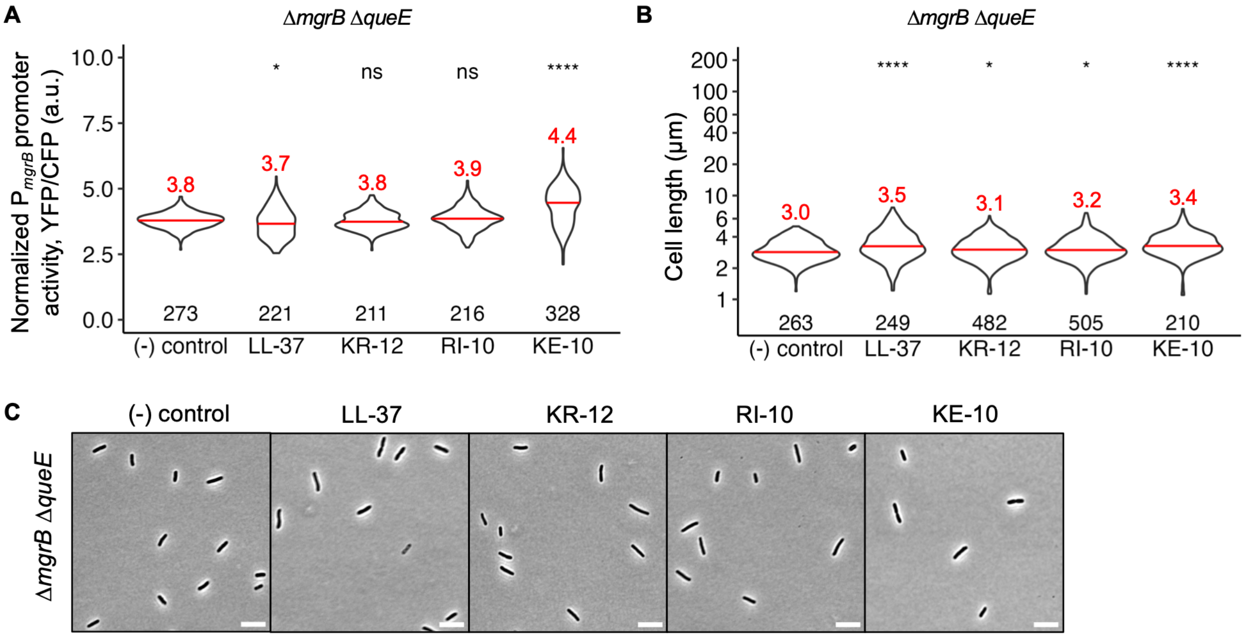

**Figure S5. PhoQP-regulated transcription and cell lengths in Δ*mgrB* Δ*queE* cells treated with sub-MIC levels of antimicrobial peptides. (A)** PhoQP-dependent promoter reporter activity, **(B)** Quantification of cell lengths, and **(C)** Representative phase-contrast micrographs of Δ*mgrB* Δ*queE* (JNC21) cells grown in the presence of sub-MIC levels of the antimicrobial peptides LL-37 (1.25 μM), KR-12 (5 μM), RI-10 (5 μM), and KE-10 (5 μM) in supplemented MinA with 0.1 mM $MgSO_4$. All data and images shown in the above panels A-C were obtained from three biological replicates; mean and median values are shown in red text and red bars, respectively. The number of cells analyzed is indicated for each sample. Significance is determined by Shapiro-Wilk test: ****P ≤ 0.0001, ***P ≤ 0.001, **P ≤ 0.01, *P ≤ 0.05, and “ns” = P > 0.05.
